# Identification of salivary and plasma biomarkers for obesity in children by non-targeted metabolomic analysis

**DOI:** 10.1101/371815

**Authors:** J. Max Goodson, Marcus Hardt, Mor-Li Hartman, Fabian Schulte, Mary Tavares, Al-Sabiha Mutawa, Jitendra Ariga, Pramod Soparkar, Jawad Behbehani, Kazem Behbehani, Mohammed Razzaque

**Affiliations:** Department of Applied Oral Sciences, the Forsyth Research Institute, Cambridge, Massachusetts, United States of America; Ministry of Health, Kuwait City, Kuwait; Kuwait School Health Program, Kuwait City, Kuwait; Faculty of Dentistry, Kuwait University, Kuwait City; The Dasman Diabetes Institute, Kuwait City, Kuwait

**Author notes:** Current address: Lake Erie College of Osteopathic Medicine, 1858 West Grandview Boulevard, Erie, PA 16509, USA. Corresponding author (JMG).

## Abstract

Chemical compounds in the saliva most likely to be associated with obesity are identified in a metabolomic analysis of paired whole saliva and plasma samples from 68 children (10-year old) who have also been evaluated for their gingival redness.

**Results:** Through metabolomic analysis119 compounds were found only in saliva, 210 only in plasma and 126 in both. The most common plasma metabolites were lipids. The most common saliva metabolites were peptides. Amino acids and their metabolites were common in both samples.

Surrogate indicators were identified by computing correlations between saliva and plasma. 29 of the 126 found in both saliva and plasma had significant positive correlation and only 4 of those (urate, creatinine, pipecolate and hydroxyproline) were associated with obesity as potential surrogate biomarkers. The uremic toxin N1-Methyl-2-pyridone-5-carboxamide (2PY) was also elevated in the saliva of obese children. 21 biochemicals were elevated in both obesity and gingivitis suggesting that some biochemical pathways for gingivitis and obesity are shared.

Of the metabolites found only in saliva, 35 were associated with obesity (p<0.01). The most significant was phosphate. Saliva had 53 dipeptides, seven of which were associated with obesity. Two volatile amines (putrescine and cadaverine) and their amino acid precursors (ornithine and lysine) were also associated with obesity.

Of the metabolites found only in plasma, 64 were associated with obesity (p<0.01). Significant increases in branched-chain and aromatic amino acids, significant reductions in serotonin, serine, and glycine and increased androgen metabolism are changes observed in 10-year old obese children that have also been reported with adult patients having type II diabetes.

**Conclusions:** Salivary urate may be a valuable measure of metabolic disease particularly in association with fructose consumption. Elevated salivary creatinine levels and presence of 2PY suggests a possible association of obesity with developing kidney disease in obese children. Though not a surrogate variable, salivary phosphate, (AUROC= 0.8) could also be an important indicator of obesity. Cadaverine, putrescine and dipeptides are also likely oral bacterial products that could help define the metabolic pathways responsible for obesity. The results of this study indicate that salivary metabolites can be important indicators of developing metabolic disease in children and provide a biochemical signature in the pathways that lead to obesity.

## Introduction

Metabolites found in saliva hold importance in providing both systemic disease insight and diagnostic potentiality. In a study of childhood obesity, we have evaluated saliva and plasma obtained from 10-year old children for metabolomic biomarkers indicating disease. The search for markers of early weight gain in children has lead us to the evaluation of children of differing body weight to identify obesity-related metabolites. From these samples we have explored the identification of potential saliva surrogate variables by their occurrence in both saliva and plasma as well as the identification of compounds of known identity (biochemicals) that are unique to saliva and not found in plasma and those unique to plasma and not found in saliva. The objective of this study was to identify obesity-related metabolites in saliva and plasma samples from children.

## Materials and methods

### Patients

Whole saliva and plasma samples were obtained from sixty-eight children of both sexes (43 boys and 25 girls) between 7 and 15 years of age. Subjects were recruited by advertisement both in Cambridge, Massachusetts and Portland, Maine from February 2011 to September 2011. The research protocol was reviewed and approved by the Forsyth Institutional Review Board. Both parental informed consent and the child’s assent were obtained prior to their enrollment.

### Saliva and blood sample collection

In each case, saliva and blood samples were collected under fasting conditions (in the morning before breakfast or at least 3 hours after a meal). A sample of 5 ml blood and 3 ml saliva were collected from each. Each child rinsed their mouth with 15 mL of water before saliva collection. Whole saliva (approximately 3 mL) was collected in labeled, sterile, 15-mL plastic screw-top centrifuge tubes (#430791, Corning Incorporated Life Sciences, Tewksbury, Massachusetts) by unstimulated drooling using standard techniques [1].

Saliva samples maintained on ice were rapidly transported to a laboratory where they were centrifuged at 2,800 RPM for 20 minutes to remove particulate debris and exfoliated mucosal cells. Supernatant saliva was transferred to screw-cap, 1-mL, 2D-barcoded storage tubes (Matrix^®^ 2D Barcoded ScrewTop^®^ storage tubes, Thermo Fisher Scientific Inc., Hudson, New Hampshire) and read by a barcode reader (Thermo Scientific VisionMate^®^ ST Barcode Reader, Thermo Fisher Scientific Inc., Hudson, New Hampshire), by which participant number was related to sample number.

Blood was taken by a phlebotomist from the median cubital artery (BD Hemogard plus, K2 BD367863, lavender cap), centrifuged 10 minutes at 2,000 x g in a refrigerated centrifuge. Both saliva and plasma samples were stored at −80°C until assayed.

### Clinical Assessments

All clinical evaluations were conducted by trained examiners. Height was measured by a stadiometer and weight was measured by a calibrated bathroom scale. Blood pressure and heart rate were measured by an automated cuff reading after the children had sat quietly for 10 minutes. Dental measurements were obtained using a portable pediatric dental chair, intraoral light and mouth mirror. Gingivitis was evaluated by counting the number of red gingival sites (mesial, buccal, distal, and lingual of each tooth) by dental personnel. Dental decay was evaluated by counting the number of teeth (both deciduous and permanent) that had visible decay or were filled. Neither dental probes nor radiographs were used in this study.

### Metabolomic analysis

120 μL aliquots of saliva supernatants and of plasma samples from each participant were assayed. Relative amounts of each metabolite were obtained by integrating peaks detected on a untargeted metabolic profiling platform (Metabolon®, Durham, North Carolina) employing high performance liquid chromatography/tandem mass spectrometry (HPLC-MS/MS and gas chromatography-mass spectrometry (GC-MS) for volatile species [2]. Compounds were identified by matching chromatographic retention times and mass spectral fragmentation signatures with reference library data created from authentic standards.

### Data analysis

All 455 biochemicals identified in saliva, in plasma or in both were subjected to three levels of analyses. The first analysis was to determine whether a saliva biochemical might be a plasma surrogate. The second analysis was to determine whether a biochemical predicts obesity. The third analysis was to determine a biochemical predicts gingivitis, hypertension or dental caries. Patient information and the quantitation data of this study is provided in the supplementary tables. S1 Table contains the clinical categorical data. S2 Table contains the data for saliva and S3 Table the data for plasma. Only those biochemicals with p-values for obesity ≤ 0.01 were retained in tables within the main text.

At the primary level, an analysis was conducted to determine the correlation between saliva levels and plasma levels identifying those biochemicals in saliva that may be directly related to plasma. This analysis, if positive and significant would indicate whether a positive distribution coefficient between the plasma and whole saliva compartments exists. If negative or insignificant, there is likely no relationship between saliva and plasma levels so that these variables are unlikely surrogate variable candidates. This assumes, however, that the concentration in both saliva and plasma are within the measurement sensitivity of the mass spectrometric method.

For the secondary level, the ability to predict whether a subject is obese is tested by non-parametric statistics (Mann-Whitney U test). If this measure was significant, the biochemical was identified as a potential biomarker for obesity. If not, it was discarded. This analysis was augmented by determining the area under the receiver operating curve (AUROC). This measure is a summary measure of the accuracy of a quantitative diagnostic test which may be interpreted as the average sensitivity for all possible values of specificity [3]. Values for AUROC were determined from metabolomic data using MetaboAnalyst 3.0 software (http://www.metaboanalyst.ca/).

For the third level, the ability to predict whether a subject has gingivitis, dental caries or high blood pressure was tested by non-parametric statistics. The results indicate whether any of the biochemicals (either in saliva or plasma) were potential biomarkers for these disease conditions and whether they conflicted with the identification of obesity.

<u>Clinical assessment</u>: A child with obesity was defined as having ≥95^th^ percentile in body mass index (BMI) (https://nccd.cdc.gov/dnpabmi/Calculator.aspx). A child with gingivitis was defined as those with more than the median percentage of red gingival sites (10.6% of sites red). Children with dental decay were defined as those with any visible decay on any tooth. Hypertension was defined as having systolic blood pressure ≥ 130 mmHg.

## Results and discussion

### Patient demographics

Of the 68 children evaluated (Table 1), 22 were obese (16 male, 6 female) and 46 were not obese (6 male, 40 female) with a mean age of 10.7± 0.2 years. The mean BMI, waist circumference, body weight, gingival redness, height, age and systolic blood pressure of obese children were significantly greater in obese children. No difference was seen in diastolic blood pressure or dental decay.

**Table 1.** Anthropomorphic and clinical characteristics of the study population grouped by obesity. Listed values are mean ± standard error. p-values refer to an analysis by t-test.

| Parameter | Overall | Obese | Not Obese | p-Value |
| --- | --- | --- | --- | --- |
| Number of subjects | 68 | 22 | 46 |  |
| Body mass index (kg/m<sup>2</sup>) | 22.2 ± 0.9 | 29.6 ± 1.5 | 18.7 ± 0.6 | 0.0000003 |
| Waist (cm) | 74.4 ± 2.3 | 93. ± 4.0 | 63.5 ± 1.6 | 0.000001 |
| Weight (kg) | 50.9 ± 3.1 | 73.6 ± 6.4 | 38.6 ± 1.9 | 0.00004 |
| Gingival redness (%) | 16.6 ± 2.4 | 29.9 ± 5.9 | 6.7 ± 1.4 | 0.003 |
| Height (cm) | 148.1 ± 1.6 | 155.2 ± 3.3 | 144.8 ± 1.6 | 0.008 |
| Age (years) | 10.7 ± 0.2 | 11.5 ± 0.4 | 10.3 ± 0.2 | 0.02 |
| Systolic blood pressure (mmHg) | 120.5 ± 1.3 | 124.1 ± 2.1 | 118.7 ± 1.6 | 0.05 |
| Diastolic blood pressure (mmHg) | 68.5 ± 0.9 | 69.8 ± 1.8 | 68.5 ± 1.0 | 0.36 |
| Dental decay (%) | 1.9 ± 0.6 | 2.3 ± 1.1 | 0.0 ± 0.7 | 0.72 |

#### Distribution of biochemicals

A combined number of 455 unique biochemicals were identified from the plasma and centrifuged saliva supernatant samples (Figure 1). 119 (26%) were detected only in saliva, 210 (46%) were found only in plasma and 126 (28%) were found in both plasma and saliva.

**Figure 1.**
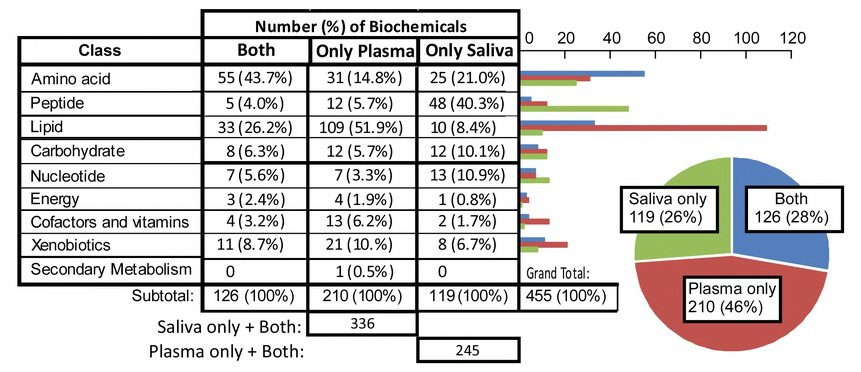
The number and percentage of biochemicals found in saliva only, in plasma only and in both by metabolomic analysis of plasma and saliva samples from the same individuals.

The majority of biochemicals that are common between saliva and plasma were amino acids and their metabolites (Figure 1). In saliva, peptides were the most commonly found biochemicals. In plasma, the most common were lipids. 109 (77%) lipids species found in plasma were not detected in saliva and 48 (91%) of the peptides found in saliva were not detected in plasma.

Analysis by obesity, gingivitis, caries and hypertension demonstrated that obesity had the greatest representation by both salivary and plasma metabolites (Figure 2). Obesity was significantly (p ≤ 0.01) associated with 39 biochemicals by both salivary analysis (Table 3) and 64 by plasma analysis (Table 4).

**Figure 2.**
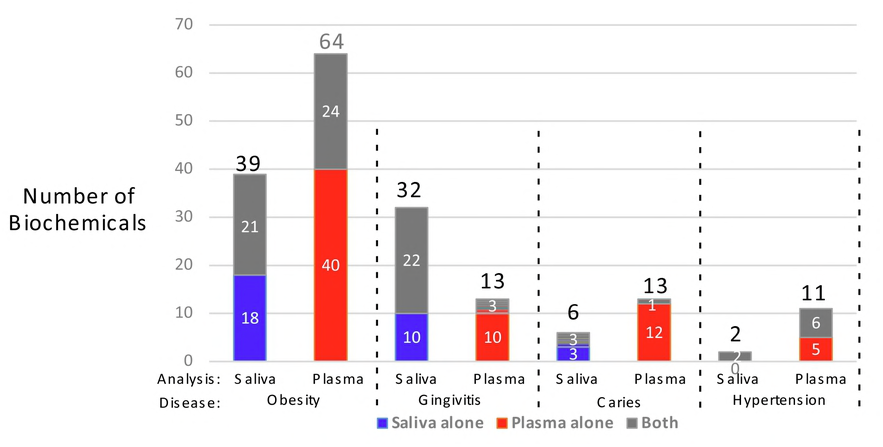
Several saliva and plasma-based biochemicals significantly (p≤ 0.01) associated with obesity, gingivitis, caries or hypertension.

### Surrogate Biochemicals

Analysis of saliva and plasma identified 29 biochemicals with significant positive correlation (Table 2). The saliva levels of the top four (pipecolate, creatinine, trans-4-hydroxyproline and urate) were also significantly associated with obesity. Regression analyses of these biochemicals (Figure 3) reveals that these four metabolites were significantly correlated.

**Figure 3.**
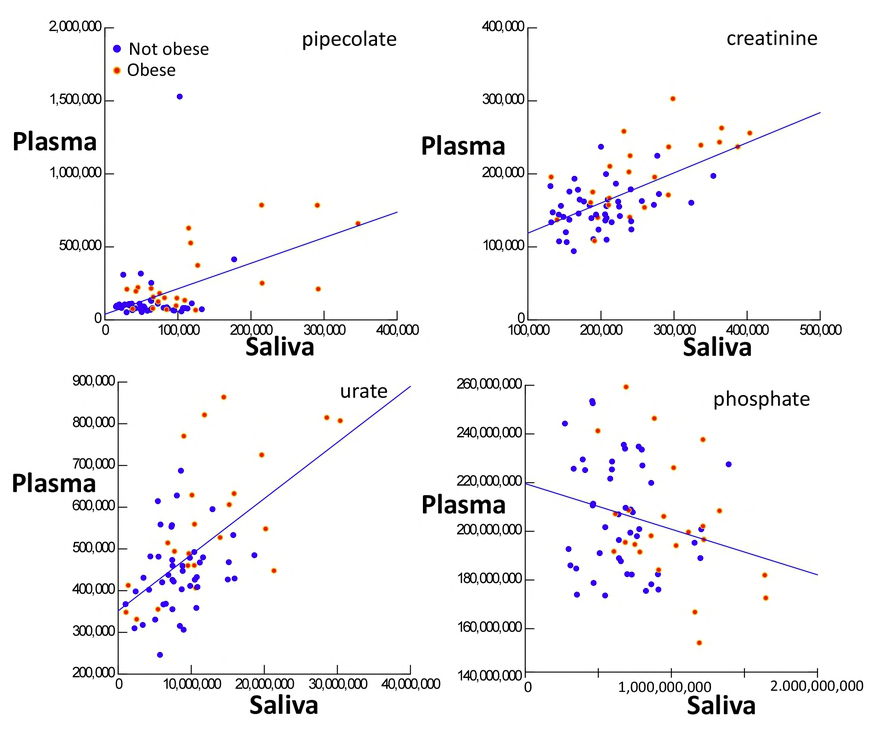
Correlation between plasma and saliva mass spectrometric values with 95% confidence intervals for the four salivary biochemicals that were also significantly related to obesity.

**Pipecolate** is a metabolite of lysine found in human urine, plasma, saliva and CSF that likely comes either from food eaten or from bacterial metabolism. In our study, significant association was only found in saliva. Recent studies suggest that plasma pipecolic acid, particularly the D-isomer, originates mainly from the catabolism of dietary lysine by intestinal bacteria rather than by direct food intake. In bacteria, pipecolate is involved with antibiotic synthesis for some species[4]. It is elevated in the plasma of patients with Zellweger syndrome and in chronic liver disease [5]. Pipecolate was also a significant indicator of gingivitis (Table 2).

**Table 2.** Biochemicals (n=29) with significant correlation (p ≤ 0.01) between saliva and plasma sorted in the order of the p-value for prediction of obesity p (Obesity) by saliva biochemicals. N is the number of children in which the biochemical was detected. Correlation p is the p-value of a correlation between saliva and plasma values. AUROC is the area under the ROC curve that predicts obesity as a univariate predictor. The fold value is the ratio of the mean for obese children divided by the mean for non-obese children. P-values for prediction of gingivitis, caries and hypertension in saliva and plasma are included.

| BIOCHEMICAL | SUPER_PATHWAY | N |  | Correlation | p(Obesity) |  | AUROC |  | Fold |  | p(Gingivitis) |  | p(Dental caries) |  | p(Hypertension) |  |
| --- | --- | --- | --- | --- | --- | --- | --- | --- | --- | --- | --- | --- | --- | --- | --- | --- |
|  |  | Saliva | Plasma | p | Saliva | Plasma | Saliva | Plasma | Saliva | Plasma | Saliva | Plasma | Saliva | Plasma | Saliva | Plasma |
| pipecolate | Amino acid | 67 | 68 | 0.00003 | 0.0007 | 0.0002 | 0.76 | 0.78 | 1.96 | 1.99 | 0.01 | 0.12 | 0.23 | 0.41 | 0.59 | 0.38 |
| creatinine | Amino acid | 68 | 68 | 0.00000002 | 0.002 | 0.0003 | 0.73 | 0.77 | 1.28 | 1.30 | 0.08 | 0.04 | 0.21 | 0.11 | 0.15 | 0.53 |
| trans-4-hydroxyproline | Amino acid | 65 | 68 | 0.002 | 0.003 | 0.61 | 0.71 | 0.54 | 1.66 | 1.05 | 0.07 | 0.99 | 0.34 | 0.92 | 0.96 | 0.98 |
| urate | Nucleotide | 68 | 68 | 0.0000003 | 0.01 | 0.002 | 0.68 | 0.73 | 1.49 | 1.28 | 0.12 | 0.03 | 0.07 | 0.08 | 0.04 | 0.30 |
| 1,5-anhydroglucitol (1,5-AG) | Carbohydrate | 68 | 68 | 0.0001 | 0.02 | 0.16 | 0.68 | 0.60 | 1.39 | 1.10 | 0.47 | 0.18 | 0.64 | 0.85 | 0.17 | 0.95 |
| 1,2-propanediol | Lipid | 68 | 68 | 0.00000002 | 0.03 | 0.07 | 0.66 | 0.63 | 2.77 | 6.83 | 0.13 | 0.38 | 0.34 | 0.17 | 0.49 | 0.88 |
| carnitine | Lipid | 68 | 68 | 0.004 | 0.03 | 0.01 | 0.66 | 0.69 | 1.27 | 1.08 | 0.03 | 0.77 | 0.60 | 0.64 | 0.13 | 0.38 |
| phenol sulfate | Amino acid | 68 | 68 | 0.00000002 | 0.04 | 0.22 | 0.65 | 0.59 | 1.41 | 1.16 | 0.01 | 0.90 | 0.56 | 0.83 | 0.36 | 0.73 |
| N1-Methyl-2-pyridone-5-carboxamide | Cofactors and vitamins | 67 | 55 | 0.0001 | 0.04 | 0.20 | 0.66 | 0.55 | 1.36 | 1.17 | 0.14 | 0.33 | 0.01 | 0.05 | 0.08 | 0.24 |
| theophylline | Xenobiotics | 19 | 44 | 0.003 | 0.04 | 0.35 | 0.54 | 0.57 | 1.91 | 1.49 | 0.08 | 0.13 | 0.54 |  | 0.24 | 0.98 |
| 2-hydroxybutyrate (AHB) | Amino acid | 66 | 68 | 0.002 | 0.05 | 0.52 | 0.63 | 0.55 | 1.31 | 1.03 | 0.18 | 0.45 | 0.94 | 0.23 | 0.13 | 0.26 |
| stachydrine | Xenobiotics | 64 | 68 | 0.00000002 | 0.09 | 0.50 | 0.63 | 0.55 | 1.77 | 1.70 | 0.35 | 0.66 | 0.07 |  | 0.95 | 0.89 |
| cortisone | Lipid | 61 | 68 | 0.00006 | 0.09 | 0.79 | 0.58 | 0.52 | 1.14 | 1.02 | 0.08 | 0.46 | 0.49 | 0.10 | 0.69 | 0.42 |
| p-cresol sulfate | Amino acid | 64 | 68 | 0.00000006 | 0.11 | 0.002 | 0.63 | 0.73 | 0.69 | 0.62 | 0.34 | 0.41 | 0.39 | 0.15 | 0.95 | 0.51 |
| hippurate | Xenobiotics | 54 | 68 | 0.00000002 | 0.13 | 0.86 | 0.64 | 0.51 | 1.14 | 0.81 | 0.75 | 0.39 | 0.27 | 0.17 | 1.00 | 0.30 |
| tryptophan betaine | Amino acid | 36 | 66 | 0.00000002 | 0.13 | 0.85 | 0.58 | 0.52 | 0.61 | 0.77 | 0.74 | 0.35 | 0.03 | 0.82 | 0.87 | 0.41 |
| catechol sulfate | Xenobiotics | 64 | 68 | 0.00000002 | 0.15 | 0.10 | 0.64 | 0.62 | 0.99 | 1.08 | 0.75 | 0.48 | 0.47 | 0.07 | 0.46 | 0.26 |
| caproate (6:0) | Lipid | 42 | 68 | 0.01 | 0.23 | 0.01 | 0.61 | 0.68 | 0.75 | 0.86 | 0.26 | 0.63 | 0.92 | 0.21 | 0.43 | 0.07 |
| propionylcarnitine | Lipid | 68 | 68 | 0.002 | 0.26 | 0.11 | 0.58 | 0.62 | 1.22 | 1.09 | 0.97 | 0.55 | 0.65 | 0.67 | 0.05 | 0.001 |
| theobromine | Xenobiotics | 62 | 64 | 0.00000002 | 0.30 | 0.17 | 0.57 | 0.57 | 2.02 | 1.76 | 0.09 | 0.28 | 0.07 |  | 0.29 | 0.38 |
| fructose | Carbohydrate | 65 | 68 | 0.00000002 | 0.33 | 0.49 | 0.57 | 0.55 | 1.49 | 1.18 | 0.18 | 0.60 | 0.75 | 0.88 | 0.86 | 0.78 |
| 3-hydroxybutyrate (BHBA) | Lipid | 62 | 68 | 0.00000005 | 0.40 | 0.02 | 0.55 | 0.67 | 0.83 | 0.39 | 0.06 | 0.74 | 0.41 | 0.18 | 0.99 | 0.26 |
| paraxanthine | Xenobiotics | 48 | 24 | 0.00000012 | 0.40 | 0.16 | 0.50 | 0.55 | 2.11 | 2.04 | 0.93 | 0.31 | 0.30 |  | 0.86 | 0.92 |
| isovaleryl carnitine | Amino acid | 66 | 68 | 0.0002 | 0.42 | 0.06 | 0.58 | 0.64 | 0.88 | 1.23 | 0.59 | 0.52 | 0.18 | 0.05 | 0.05 | 0.002 |
| betaine | Amino acid | 68 | 68 | 0.004 | 0.50 | 0.20 | 0.55 | 0.60 | 0.99 | 0.95 | 0.13 | 0.51 | 0.92 | 0.77 | 0.66 | 0.63 |
| 3-indoxyl sulfate | Amino acid | 56 | 68 | 0.00000002 | 0.51 | 0.09 | 0.56 | 0.63 | 1.10 | 0.80 | 0.80 | 0.26 | 0.60 | 0.67 | 0.24 | 0.30 |
| erythritol | Xenobiotics | 49 | 68 | 0.001 | 0.58 | 0.01 | 0.61 | 0.70 | 1.19 | 1.18 | 0.35 | 0.07 | 0.29 |  | 0.37 | 0.60 |
| 2-methylbutyrylcarnitine | Amino acid | 45 | 68 | 0.0003 | 0.71 | 0.06 | 0.57 | 0.64 | 0.99 | 1.15 | 0.63 | 0.20 | 0.73 | 0.41 | 0.68 | 0.06 |
| caffeine | Xenobiotics | 47 | 65 | 0.00000002 | 0.75 | 0.74 | 0.52 | 0.52 | 1.57 | 1.68 | 0.78 | 0.83 | 0.20 |  | 0.07 | 0.17 |

***Creatinine*** is a metabolite of creatinine phosphate which is released from muscle at a relatively constant rate. Blood levels of creatinine (100 mM=1 mg/dL) are clinically important since they are used to estimate glomerular filtration rate. Of the biomarkers in our study, creatinine had the highest saliva:plasma correlation measured (0.48) suggesting that saliva concentrations of this metabolite were a surrogate for plasma levels. Significantly elevated salivary levels of creatinine were observed in obese children (Table 2), which also suggests a possible association with kidney disease [6]. The data of Table 2 and Figure 3 suggest that salivary creatinine may be used as a surrogate variable (AUROC=0.73) for blood creatinine and as an indicator of developing obesity.

**Trans-4-hydroxyproline** in saliva is often associated with the enzymatic digestion of collagen in gingivitis and periodontal disease. The association found in this study, however suggests a possible modulation by obesity. Our data indicates no significant association. Although trans-4-hydroxyproline levels in saliva and plasma were positively correlated (p=0.002, Table 2), only saliva levels were indicative of obesity (p=0.003 for saliva and p=0.61 for plasma) suggesting an interaction between the inflammatory conditions of obesity and gingivitis. Plasma levels of trans-4-hydroxyproline were not significantly associated with obesity.

***Urate*** is generally recognized as a marker of adiposity [7], obesity and glomerular filtration [8]. Others have indicated that it is a marker of inflammation, endothelial dysfunction and elevated measures of metabolic syndrome [9]. Many consider it to have a causal role in the development of diabetes due to its role in the metabolism of fructose [10]. In our research, we provide evidence that salivary urate is a surrogate for plasma urate (Table 3). Our results also indicate a fair univariate prediction value (AUROC=0.68) for obesity.

**Table 3.** Biochemicals (n=21) in saliva associated with both obesity and gingivitis sorted by p-value for obesity. Compartment: S=saliva only, B=both plasma and saliva. Correlation is for compartment B only. Rank is based on minimum p-value for obesity and for gingivitis.

| BIOCHEMICAL | SUPER_PATHWAY | Obesity |  | Gingivitis |  | Compartment | Correlation |  | Rank |  |
| --- | --- | --- | --- | --- | --- | --- | --- | --- | --- | --- |
|  |  | p | AUROC | p | AUROC |  | (r) | p | Obesity | Gingivitis |
| phosphate | Energy | 0.00007 | 0.80 | 0.007 | 0.69 | B | -0.25 | 0.04 | 1 | 17 |
| 3-(4-hydroxyphenyl)lactate | Amino acid | 0.00009 | 0.79 | 0.0002 | 0.77 | B | 0.17 | 0.17 | 2 | 2 |
| thymine | Nucleotide | 0.0004 | 0.78 | 0.001 | 0.75 | S |  |  | 3 | 7 |
| allantoin | Nucleotide | 0.001 | 0.67 | 0.0003 | 0.59 | B | -0.11 | 0.54 | 4 | 3 |
| pipecolate | Amino acid | 0.001 | 0.76 | 0.008 | 0.70 | B | 0.48 | 0.00003 | 5 | 18 |
| 2-hydroxyglutarate | Lipid | 0.001 | 0.76 | 0.001 | 0.75 | B | -0.06 | 0.65 | 6 | 5 |
| fucose | Carbohydrate | 0.002 | 0.74 | 0.005 | 0.70 | S |  |  | 7 | 15 |
| 5-aminovalerate | Amino acid | 0.002 | 0.73 | 0.002 | 0.72 | S |  |  | 8 | 9 |
| alpha-hydroxyisocaproate | Amino acid | 0.002 | 0.73 | 0.013 | 0.69 | B | 0.03 | 0.87 | 9 | 20 |
| 3-phosphoglycerate | Carbohydrate | 0.003 | 0.67 | 0.0001 | 0.71 | S |  |  | 10 | 1 |
| agmatine | Amino acid | 0.003 | 0.69 | 0.002 | 0.73 | S |  |  | 11 | 8 |
| phenyllactate (PLA) | Amino acid | 0.003 | 0.72 | 0.002 | 0.72 | B | 0.05 | 0.70 | 12 | 10 |
| cadaverine | Amino acid | 0.003 | 0.70 | 0.009 | 0.72 | S |  |  | 13 | 19 |
| putrescine | Amino acid | 0.004 | 0.71 | 0.002 | 0.72 | S |  |  | 14 | 11 |
| erythronate | Carbohydrate | 0.005 | 0.63 | 0.014 | 0.54 | B | 0.26 | 0.12 | 15 | 21 |
| 5,6-dihydrouracil | Nucleotide | 0.006 | 0.71 | 0.004 | 0.70 | B | 0.02 | 0.85 | 16 | 13 |
| ornithine | Amino acid | 0.007 | 0.70 | 0.004 | 0.70 | B | -0.07 | 0.62 | 17 | 14 |
| alpha-hydroxyisovalerate | Amino acid | 0.01 | 0.70 | 0.002 | 0.72 | B | 0.17 | 0.18 | 18 | 12 |
| 3-aminoisobutyrate | Nucleotide | 0.01 | 0.74 | 0.006 | 0.76 | S |  |  | 19 | 16 |
| N-acetylputrescine | Amino acid | 0.01 | 0.68 | 0.001 | 0.74 | S |  |  | 20 | 4 |
| phenylacetate | Amino acid | 0.01 | 0.68 | 0.001 | 0.73 | B | -0.13 | 0.35 | 21 | 6 |

#### Salivary biochemicals associated with both obesity and gingivitis

Twenty-one salivary biochemicals were significantly related to both obesity and gingivitis (Table 3) suggesting that there may be common biochemical pathways between these two inflammatory diseases. Examining compartments of biochemicals in Table 2 indicates that with one exception (pipecolate), all biochemicals are either found only in saliva (Compartment=S) or do not significantly correlate between saliva and plasma. Hence, all biochemicals associated with both obesity and gingivitis were essentially restricted to saliva.

**Hydroxyphenyllactate** appears at second rank in both obesity and saliva. In mammalian biology this is a tyrosine metabolite but is also synthesized as the D-form by oral bacteria from phenolic compounds

**Allantoin** appears at fourth rank in obesity and at the third rank in gingivitis and is produced by animals, plants and bacteria. Allantoin, is a product of purine metabolism considered by some to be a measure of oxidative stress [11].

### Biochemicals found only in saliva

Many biochemical found only in saliva exhibited a strong microbial signature. Saliva contains approximately 10^7^ bacteria/ml [12]. General biochemical processes of proteolysis, amino acid decarboxylation, hydroxylation, methylation and acetylation appear to be the result of bacterial metabolism within the oral cavity. After surveying 342 biochemicals (245 found only in saliva plus the 126-29=94 in which saliva values were not significantly correlated with plasma) 35 were found to be obesity-associated (Table 4). The strongest predictive value of all salivary biochemicals measured was that of elevated **phosphate** with an AUROC of 0.80. Although phosphate is found both in saliva and plasma, the correlation was negative indicating that the salivary and blood compartments are not in direct communication. The top two biochemicals that correlate with salivary phosphate were glycerol (r=0.85) and 3-phosphoglycerate (r=0.69), suggesting that phosphate could be related to dephosphorylation of glyceryl phosphates to produce glycerol. The concentration of phosphate in saliva is approximately three times that of the concentration in plasma [13] and the transport of phosphate in salivary glands involves sodium-phosphate co-transporters [14]. The negative correlation between saliva and plasma seen in Table 2 can be considered evidence that the oral and plasma compartments for phosphate may not directly participate in the molecular exchange. Of relevance, high dietary phosphate consumption is found to be associated with higher occurrence of dental decay [15]. while increased salivary phosphate concentration has shown to be a potential biomarker for the evolvement of childhood obesity [16].

**Table 4.** Biochemicals identified as potential biomarkers for obesity found only in saliva (n=35) sorted by the ability to identify obese children (p≤ 0.01). These data include those found only in saliva (compartment=S) and those in both (compartment B) whose saliva/plasma correlation is ≤0.01. Other variables are those as defined in Table 2.

| BIOCHEMICAL | SUPER_PATHWAY | p(Obesity) | Fold | AUROC | N | Compartment | p(Gingivitis) | p(Caries) | p(Hypertension) |
| --- | --- | --- | --- | --- | --- | --- | --- | --- | --- |
| phosphate | Energy | 6.72E-05 | 1.49 | 0.80 | 68 | B | **0.007** | 0.65 | 0.10 |
| 3-(4-hydroxyphenyl)lactate | Amino acid | 9.30E-05 | 1.42 | 0.79 | 68 | B | **0.0002** | 0.58 | 0.73 |
| thymine | Nucleotide | 3.53E-04 | 1.61 | 0.78 | 66 | S | **0.001** | 0.68 | 0.50 |
| allantoin | Nucleotide | 5.13E-04 | 2.29 | 0.67 | 45 | B | **0.0003** | 0.78 | 0.73 |
| 2-hydroxyglutarate | Lipid | 1.05E-03 | 1.66 | 0.76 | 66 | B | **0.0009** | 0.57 | 0.78 |
| fucose | Carbohydrate | 1.53E-03 | 1.69 | 0.74 | 68 | S | **0.005** | 0.89 | 0.60 |
| cyclo(gly-phe) | Dipeptide | 1.61E-03 | 1.34 | 0.60 | 38 | S | 0.04 | 0.31 | 0.12 |
| leucylisoleucine | Dipeptide | 2.23E-03 | 1.37 | 0.67 | 61 | S | 0.12 | 0.52 | 0.81 |
| 5-aminovalerate | Amino acid | 2.27E-03 | 1.90 | 0.73 | 68 | S | **0.002** | 0.18 | 0.45 |
| alpha-hydroxyisocaproate | Amino acid | 2.49E-03 | 1.69 | 0.73 | 67 | B | **0.01** | 0.36 | 0.90 |
| 3-phosphoglycerate | Carbohydrate | 2.53E-03 | 1.78 | 0.67 | 52 | S | **0.0001** | 0.54 | 0.16 |
| agmatine | Amino acid | 3.04E-03 | 2.14 | 0.69 | 67 | S | **0.002** | 0.35 | 0.36 |
| phenyllactate | Amino acid | 3.32E-03 | 1.46 | 0.72 | 68 | B | **0.002** | 0.48 | 0.88 |
| glycylphenylalanine | Dipeptide | 3.38E-03 | 1.99 | 0.56 | 47 | S | 0.38 | 0.91 | 0.59 |
| cadaverine | Amine | 3.44E-03 | 2.80 | 0.70 | 65 | S | **0.009** | 0.36 | 0.52 |
| glycerol | Lipid | 3.47E-03 | 1.32 | 0.72 | 68 | B | 0.05 | 0.93 | 0.04 |
| N-acetylserine | Amino acid | 3.47E-03 | 1.36 | 0.59 | 56 | S | 0.38 | 0.97 | 0.56 |
| putrescine | Amine | 4.09E-03 | 1.88 | 0.71 | 68 | S | **0.002** | 0.38 | 0.38 |
| erythronate | Carbohydrate | 4.89E-03 | 1.54 | 0.63 | 38 | B | 0.01 | 0.45 | 0.12 |
| galactose | Carbohydrate | 5.21E-03 | 1.62 | 0.71 | 68 | S | 0.06 | 0.66 | 0.80 |
| 5,6-dihydrouracil | Nucleotide | 5.88E-03 | 1.27 | 0.71 | 68 | B | **0.004** | 0.74 | 0.92 |
| 3-methyl-2-oxovalerate | Amino acid | 6.79E-03 | 1.42 | 0.70 | 59 | B | 0.02 | 0.87 | 0.67 |
| ornithine | Amino acid | 6.88E-03 | 1.61 | 0.70 | 68 | B | **0.004** | 0.17 | 0.59 |
| leucylaspartate | Dipeptide | 7.07E-03 | 1.57 | 0.65 | 65 | S | 0.02 | 0.44 | 0.51 |
| uridine | Nucleotide | 7.15E-03 | 0.71 | 0.70 | 68 | B | 0.02 | 0.53 | 0.89 |
| valylphenylalanine | Dipeptide | 8.31E-03 | 1.36 | 0.63 | 41 | S | 0.08 | 0.26 | 0.92 |
| leucylserine | Dipeptide | 9.00E-03 | 1.51 | 0.69 | 68 | S | 0.04 | 0.35 | 0.10 |
| 2-hydroxy-3-methylvalerate | Amino acid | 9.03E-03 | 1.61 | 0.68 | 61 | B | 0.1369 | 0.67 | 0.90 |
| alpha-hydroxyisovalerate | Amino acid | 1.00E-02 | 1.51 | 0.70 | 67 | B | **0.002** | 0.62 | 0.91 |
| lysine | Amino acid | 1.05E-02 | 1.57 | 0.69 | 68 | B | 0.02 | 0.87 | 0.17 |
| 3-aminoisobutyrate | Nucleotide | 1.09E-02 | 1.49 | 0.74 | 53 | S | **0.006** | **0.01** | 0.77 |
| beta-alanine | Amino acid | 1.32E-02 | 1.71 | 0.50 | 44 | S | 0.02 | **0.01** | 0.54 |
| N-acetylputrescine | Amino acid | 1.35E-02 | 1.41 | 0.68 | 68 | S | **0.0006** | 0.60 | 0.33 |
| phenylacetate | Amino acid | 1.35E-02 | 1.92 | 0.68 | 68 | B | **0.0009** | 0.41 | 0.14 |
| leucylleucine | Peptide | 1.40E-02 | 1.39 | 0.68 | 68 | B | 0.11 | 0.43 | 0.16 |

Saliva biochemicals significantly associated with obesity (n=35) are listed in Table 4. Of these obesity-associated biochemicals, 21 also were significantly increased in children with gingivitis. Of the 126 biochemicals found both in saliva and plasma, 35+4=39 were found that significantly (p<0.01) associated with obesity and 12 of these were also associated with gingivitis (Table 3). Of these, pipecolate, creatinine and urate were found not only to be correlated between saliva and plasma but also to be significantly related to obesity for both saliva and plasma. Of these three biochemicals, pipecolate was also significantly related to gingivitis. Creatinine and urate, however, were not significantly associated with gingivitis, dental caries or hypertension and exhibited significant association by linear regression (Figure 3).

Saliva levels of trans-4-hydroxyproline, a recognized marker for collagen break-down, was significantly associated with obesity in saliva samples (p=0.003) but not in plasma samples (p=0.61). The p-value for gingivitis where it has been proposed as a periodontal disease marker [17] was of border line significance (p=0.07, File S3 Data).

Of the 65 peptides found in this analysis (combined salivary and plasma peptides), 54 were dipeptides (83%). Most (49, 75%) were found in saliva. Of the 152 lipids found in this analysis 109 were found only in plasma and 10 were found in saliva alone. From these observations it is clear that saliva has relatively few lipids and an extraordinary number of dipeptides.

Based on a significance cut-off of p≤0.01, there were 35 biochemicals in saliva that identified obese children (Table 4). The most sensitive biomarker in saliva was phosphate which was 1.49 times higher in obese children with an AUROC of 0.80. The most common chemical group were amino acids (19) followed by peptides (7), nucleotides (6), carbohydrates (4) and lipids (2).

Of the 48 peptides found in saliva, 45 were dipeptides and none of these were found in plasma. Six peptides were significantly related to obesity including **cyclo(gly-phe)**, **leucylisoleucine**, **glycylphenylalanine**, **leucylaspartate**, **valylphenylalanine** and **leucylserine**. Although we cannot rule out the possibility that these have been produced by the salivary gland, it seems likely that these could be indicative of proteolytic salivary bacteria.

Another group of obesity-associated variables likely of bacterial origin are the biogenic amines **cadaverine** and **putrescine** with their precursors **lysine** and **ornithine**. These volatile amines are commonly found in strong cheeses as a result of bacterial decarboxylation.

Salivary levels of cadaverine and putrescine found only in saliva were elevated in obese children (Table 3). The amino acid precursors **lysine** and **ornithine** were also found to be significantly associated with obesity both in saliva and plasma (Table 2). All four of these compounds were significantly (≤ 0.01) related to obesity and all but lysine related to gingivitis as well. Lysine decarboxylase, an enzyme that can produce cadaverine has been shown to be associated with gingival inflammation [18], denture malodor [19] and halitosis [20].

**Dipeptides** in children’s saliva, 45 dipeptides were found only in saliva. These would seem likely to represent products of bacterial metabolism since they are not found in plasma and would not be expected to occur in the diet of these children. Dipeptides have also been reported in the saliva of adults [21]. Several dipeptides were significantly increased with obesity suggesting an accentuation of proteolytic activity in the oral environment indicating increased proteolysis with increasing BMI. The 6 peptides found only in saliva that were significantly related to obesity included **cyclo(gly-phe)**, **leucylisoleucine**, **glycylphenylalanine**, **leucylaspartate**, **valylphenylalanine** and **leucylserine**. None of these dipeptides were significantly related to gingivitis, caries or hypertension. The importance of dipeptides, for the nutrition of bacterial species has been demonstrated for the intraoral species *P. gingivalis, P. intermedia, P. nigrescens and F. nucleatum*. [22] and there is reason to expect it may occur in other species as well.

Creation of dipeptides by bacterial action has been for example reported for the *P. gingivalis* dipeptidyl-peptidase IV (DPP-IV) [23]. These peptides are of the form H_2_N-Xaa-Pro. Only three salivary peptides were of that configuration (Pro-Pro, Gly-Pro and Lys-Pro) and none were particularly predictive of obesity (p>0.3)). The peptide most predictive (p=0.002) of obesity was cyclo(gly-phe) (Table 3).

### Biochemicals found only in plasma

**Serotonin** reduction in plasma was highly significant in obese children. Depression and antidepressant drugs are both associated with increased risk of obesity presumably due to a reduction in serotonin or the serotonin transporter [24, 25].

Reduced serotonin has been found in cord blood where this reduction was associated with rapid postnatal weight gain [26]. The understanding of serotonin function has been broadly expanded to include obesity and diabetes [25].

**Saturated lipids** (**caproate, heptanoate caprylate, pelargonate, caprate, laurate, myristate, nonadecoanoate**) in plasma were all significantly reduced in obesity (Tables 4 and 5). This observation differs from that usually reported for adults. Adult obesity and type 2 diabetes is generally associated with an increase in plasma free fatty acids [27].

**Table 5.** Biochemicals identified as potential biomarkers only in plasma (n=64) sorted by the ability to identify obese children (Obesity p≤ 0.01). Compartment are B=both saliva and plasma, P= plasma alone. Other variables are those described in Table 2 legend.

| BIOCHEMICAL | SUPER_PATHWAY | Obesity p | Fold | AUROC | N | Compartment | Gingivitis-p | Caries-p | Hyper-tension-p |
| --- | --- | --- | --- | --- | --- | --- | --- | --- | --- |
| serine | Amino acid | 0.000005 | 0.79 | 0.84 | 68 | B | 0.02 | 0.03 | 0.04 |
| serotonin | Amino acid | 0.00003 | 0.45 | 0.84 | 66 | P | 0.02 | 0.84 | 0.46 |
| 3-(4-hydroxyphenyl)lactate | Amino acid | 0.00006 | 1.32 | 0.80 | 68 | B | 0.21 | 0.80 | 0.0003 |
| tyrosine | Amino acid | 0.0001 | 1.20 | 0.79 | 68 | B | 0.43 | 0.90 | 0.01 |
| gamma-glutamylphenylalanine | Peptide | 0.0001 | 1.25 | 0.78 | 68 | P | 0.06 | 0.76 | 0.43 |
| pipecolate | Amino acid | 0.0002 | 1.99 | 0.78 | 68 | B | 0.12 | 0.41 | 0.38 |
| lactate | Carbohydrate | 0.0002 | 1.33 | 0.78 | 68 | B | 0.54 | 0.96 | 0.15 |
| piperine | Xenobiotics | 0.0002 | 1.72 | 0.78 | 67 | P | 0.40 | 0.27 | 0.99 |
| iminodiacetate | Xenobiotics | 0.0002 | 1.28 | 0.77 | 68 | P | 0.05 | 0.59 | 0.81 |
| creatinine | Amino acid | 0.0003 | 1.30 | 0.77 | 68 | B | 0.04 | 0.11 | 0.53 |
| valine | Amino acid | 0.0003 | 1.15 | 0.77 | 68 | B | 0.19 | 0.43 | 0.18 |
| caprate (10:0) | Lipid | 0.0006 | 0.75 | 0.76 | 68 | P | 0.09 | 0.45 | 0.43 |
| 4-androsten-3beta,17beta-diol disulfate 1 | Lipid | 0.0008 | 2.18 | 0.75 | 68 | P | 0.06 | 0.26 | 0.07 |
| pseudouridine | Nucleotide | 0.0009 | 1.12 | 0.75 | 68 | B | 0.003 | 0.20 | 0.21 |
| 4-androsten-3beta,17beta-diol disulfate 2 | Lipid | 0.001 | 1.67 | 0.74 | 68 | P | 0.12 | 0.54 | 0.06 |
| phenylacetate | Amino acid | 0.001 | 0.54 | 0.74 | 52 | B | 0.02 | 0.42 | 0.82 |
| adenosine 5'-monophosphate | Nucleotide | 0.001 | 0.57 | 0.75 | 67 | P | 0.01 | 0.80 | 0.23 |
| indolelactate | Amino acid | 0.001 | 1.34 | 0.74 | 68 | B | 0.23 | 0.93 | 0.24 |
| L-urobilin | Cofactors and vitamins | 0.002 | 0.20 | 0.65 | 42 | P | 0.61 | 0.43 | 0.22 |
| p-cresol sulfate | Amino acid | 0.002 | 0.62 | 0.73 | 68 | B | 0.41 | 0.15 | 0.51 |
| EDTA | Xenobiotics | 0.002 | 1.12 | 0.73 | 68 | P | 0.13 | 0.16 | 0.06 |
| urate | Nucleotide | 0.002 | 1.28 | 0.73 | 68 | B | 0.03 | 0.08 | 0.30 |
| isoleucine | Amino acid | 0.002 | 1.15 | 0.73 | 68 | B | 0.19 | 0.50 | 0.56 |
| leucine | Amino acid | 0.002 | 1.13 | 0.73 | 68 | B | 0.21 | 0.36 | 0.27 |
| 10-undecenoate (11:1n1) | Lipid | 0.002 | 0.66 | 0.73 | 68 | P | 0.12 | 1.00 | 0.87 |
| mannose | Carbohydrate | 0.003 | 1.15 | 0.72 | 68 | P | 0.20 | 0.69 | 0.60 |
| pyruvate | Carbohydrate | 0.003 | 1.33 | 0.72 | 68 | P | 0.39 | 0.77 | 0.42 |
| pyrophosphate | Energy | 0.003 | 1.38 | 0.72 | 68 | P | 0.72 | 0.77 | 0.41 |
| heptanoate (7:0) | Lipid | 0.003 | 0.83 | 0.72 | 68 | P | 0.36 | 0.11 | 0.03 |
| gamma-glutamyltyrosine | Peptide | 0.003 | 1.22 | 0.72 | 68 | P | 0.12 | 0.48 | 0.01 |
| pelargonate (9:0) | Lipid | 0.004 | 0.85 | 0.71 | 68 | P | 0.11 | 0.21 | 0.11 |
| laurate (12:0) | Lipid | 0.004 | 0.80 | 0.71 | 68 | P | 0.57 | 0.45 | 0.18 |
| phenylacetylglutamine | Amino acid | 0.005 | 0.64 | 0.71 | 68 | P | 0.44 | 0.48 | 0.74 |
| 7-alpha-hydroxy-3-oxo-4-cholestenoate | Lipid | 0.005 | 1.21 | 0.71 | 68 | P | 0.80 | 0.53 | 0.02 |
| hypoxanthine | Nucleotide | 0.005 | 1.44 | 0.71 | 68 | B | 0.39 | 0.31 | 0.95 |
| glutaryl carnitine | Amino acid | 0.005 | 1.20 | 0.71 | 68 | P | 0.73 | 0.70 | 0.18 |
| gamma-glutamylvaline | Peptide | 0.006 | 1.20 | 0.70 | 68 | P | 0.05 | 0.59 | 0.98 |
| gamma-glutamylisoleucine | Peptide | 0.006 | 1.31 | 0.70 | 68 | P | 0.41 | 0.50 | 0.94 |
| 15-methylpalmitate | Lipid | 0.006 | 0.81 | 0.70 | 68 | P | 0.22 | 0.45 | 0.88 |
| [H]HWESASLLR[OH] | Peptide | 0.006 | 1.39 | 0.70 | 68 | P | 0.43 | 0.99 | 0.26 |
| cysteine | Amino acid | 0.007 | 1.19 | 0.70 | 68 | B | 0.15 | 0.19 | 0.03 |
| glycine | Amino acid | 0.007 | 0.85 | 0.70 | 68 | B | 0.40 | 0.20 | 0.35 |
| erythritol | Xenobiotics | 0.007 | 1.18 | 0.70 | 68 | B | 0.07 |  | 0.60 |
| cystine | Amino acid | 0.007 | 1.36 | 0.61 | 52 | P | 0.01 | 0.55 | 0.25 |
| 21-hydroxypregnenolone disulfate | Lipid | 0.008 | 1.32 | 0.67 | 62 | P | 0.30 | 0.23 | 0.09 |
| 7-methylxanthine | Xenobiotics | 0.008 | 1.56 | 0.57 | 49 | P | 0.01 | 0.19 | 0.62 |
| N-acetyl-beta-alanine | Amino acid | 0.008 | 1.23 | 0.70 | 68 | P | 0.41 | 0.39 | 0.15 |
| acetylphosphate | Energy | 0.008 | 1.12 | 0.70 | 68 | P | 0.48 | 0.07 | 0.14 |
| S-methylcysteine | Amino acid | 0.009 | 1.49 | 0.64 | 53 | P | 0.96 | 0.29 | 0.86 |
| phenylalanine | Amino acid | 0.009 | 1.12 | 0.69 | 68 | B | 0.70 | 1.00 | 0.13 |
| caprylate (8:0) | Lipid | 0.009 | 0.81 | 0.69 | 68 | B | 0.40 | 0.25 | 0.29 |
| 3-methylxanthine | Xenobiotics | 0.009 | 1.61 | 0.53 | 32 | P | 0.21 | 0.17 | 0.19 |
| alpha-hydroxyisovalerate | Amino acid | 0.009 | 1.33 | 0.69 | 68 | B | 0.78 | 0.34 | 0.62 |
| myristate (14:0) | Lipid | 0.01 | 0.79 | 0.69 | 68 | P | 0.68 | 0.75 | 0.65 |
| arabitol | Carbohydrate | 0.01 | 1.22 | 0.70 | 67 | P | 0.45 | 0.30 | 0.05 |
| phenyllactate | Amino acid | 0.01 | 1.23 | 0.70 | 67 | B | 0.09 | 0.11 | 0.25 |
| 4-hydroxyhippurate | Xenobiotics | 0.01 | 1.71 | 0.64 | 61 | P | 0.51 | 0.48 | 0.51 |
| 5alpha-androstan-3beta,17beta-diol disulfate | Lipid | 0.01 | 1.84 | 0.75 | 47 | P | 0.03 | 0.46 | 0.88 |
| carnitine | Lipid | 0.01 | 1.08 | 0.69 | 68 | B | 0.77 | 0.64 | 0.38 |
| caproate (6:0) | Lipid | 0.01 | 0.86 | 0.68 | 68 | B | 0.63 | 0.21 | 0.07 |
| nonadecanoate (19:0) | Lipid | 0.01 | 0.71 | 0.68 | 68 | P | 0.11 | 0.45 | 0.40 |
| gamma-glutamylleucine | Peptide | 0.01 | 1.18 | 0.68 | 68 | P | 0.43 | 0.71 | 0.75 |
| laurylcarnitine | Lipid | 0.01 | 0.65 | 0.71 | 66 | P | 0.20 | 0.94 | 0.80 |
| beta-sitosterol | Lipid | 0.01 | 1.34 | 0.55 | 33 | P | 0.10 | 0.47 | 0.96 |

Plasma biochemicals significantly related to obesity (n=64) are listed in Table 5.

Based on a significance cut-off of p≤0.01, there were 64 biochemicals in plasma that identified obese children (Table 4). The most sensitive biomarkers in plasma were serine (0.79-fold reduction) and serotonin (0.45 fold reduction) both with an AUROC of 0.84.

The most significant change in biochemicals associated with obesity in plasma was reduction in serotonin (Table 5) with obesity.

**Gamma-glutamylphenylalanine** increase with obesity suggests an increase in gamma-glutamyl transferase (GGT, E.C. 2.3.3.2) activity. GGT has long been used to evaluate liver disease and is also associated with diabetes [28]. The third strongest biochemical, **piperine** is one of the alkaloids that gives pepper its pungency.

**Amino acids and amino acid derivatives** were common (n=22) in the biochemicals that predict obesity. Three branched chain amino acids (**valine, isoleucine, leucine**) and two aromatic amino acids (**tyrosine, and phenylalanine**) significantly increased in plasma of obese children. Longitudinal analysis of adults in the Framingham Heart Study [29], demonstrate that elevated plasma levels of these five amino acids stimulate insulin release and predict future development of diabetes up to 12 years before the onset of diabetes. Two amino acids (**serine and glycine**) significantly decreased with obesity. The reduction in serine with obesity was the most powerfully associated biochemical change in plasma samples (p=0.00001, AUC=0.84)) which has also been reported by other investigators [30].

### Biochemicals of special interest

The uremic toxin **N1-Methyl-2-pyridone-5-carboxamide** (2PY), found in both saliva and plasma increased in obese children (p= 0.04 AUROC=0.66 in saliva). **1,5 anhydroglucitol (1,5-AG)** is a naturally occurring non-metabolizable monosaccharide found in most foods and was detected in both saliva and plasma. Because of its ability to compete with glucose for renal excretion it has been developed as an assay for hyperglycemia in adults. In our study, both saliva and plasma levels were increased in obese children but not to the significance level of predicting obesity with p≤ 0.01. **Metformin** is often used to treat type 2 diabetes in children. It was found in both saliva and plasma of one obese child. **α-hydroxy butyrate** has been proposed as an early plasma biomarker of insulin resistance in adults [31]. It was found in all plasma and all but two saliva samples, but the prediction of obesity from saliva samples was borderline (p≤0.05, AUROC=0.63) and prediction from plasma samples (p=0.52, AUROC=0.55) far from significance. This could suggest that this biomarker may be in the development phase for 10-year old children. **Glucose,** a common clinical measure of diabetic effects in adults was measured in the saliva of all samples, but it was not included as a proposed biomarker for obesity because few obese children developed hyperglycemia. From other studies [32] we know that the concentration of glucose (86.3 mg/dL) in plasma is about two orders of magnitude greater than in saliva (0.11 mg/dL) and few obese children of 10 years age had elevated salivary glucose. **Melatonin**, suggested by many to occur in saliva [33] was not found in blood or saliva by mass spectrometry in our studies.

**Limitations of this study** include the method of non-targeted metabolomic analysis. This method does not provide an estimate of assay sensitivity or absolute concentrations for individual biochemicals so that there is always an unknown factor associated with this analysis. This is not a limitation of mass spectrometry but a limitation of the non-targeted method which provided only relative amounts of 455 biochemicals but no absolute concentration values. Having identified a subset of 35+4=39 potentially important salivary biochemicals, however, makes future targeted analysis of promising biochemicals more feasible by eliminating biochemicals that do not appear to be related to obesity.

Another limitation is the problem of multiple analyses. Only those biochemicals with p values for obesity ≤ 0.01 were retained in text tables. Since the Bonferroni corrected p value of 0.05 for 455 comparisons is 0.0001, true statistical significance occurred only for pipecolate (Table 2), phosphate and 3-4 hydroxyphenyl lactate (Table 3) and serine (Table 5). The purpose of this analysis is to identify those biochemicals in the saliva which are most likely to be associated with obesity and least associated with other common systemic disease conditions and not to prove clinical diagnostic capability. True analysis of diagnostic potential will require a targeted approach with a much smaller number of biochemicals tested across a larger cohort.

## Conclusions

Saliva is capable of providing considerable information related to developing obesity in children. Since it does not appear, however, to be adapted to the estimation of many fat-soluble metabolites in plasma. Most metabolites found only in saliva are likely derived from bacterial metabolism. Salivary urate may be a useful estimator of fructose consumption and risk of metabolic syndrome. Increased salivary creatinine and 2PY suggest possible early renal disease in obese children. Plasma levels of many metabolites in obese (not diabetic) children differed in the same direction as adults with type II diabetes. Finally, salivary phosphate content might reflect the evolvement of childhood obesity. The results of our non-targeted metabolomic analysis form the basis for further targeted studies to precisely identify the roles and regulation of identified factors in obesity and metabolic diseases, in general.

## Supporting information

**S1 Table.** Subject number and binary values describing sex, obesity, gingivitis high blood pressure and dental caries.

**S2 Table.** Original scale mass spectral values for all detected biochemicals in saliva.

**S3 Table.** Original scale mass spectral values for all detected biochemicals in plasma.

